# Dosage-dependent disruption of corticofugal neuron development by Fezf2 loss

**DOI:** 10.64898/2026.10.21.760692

**Authors:** Md. Jubayer Hossain, Sheikh Naem Islam Abhi, Md. Mahfuzul Islam, Muhibullah Shahjahan, Umme Marzia Tasnim, Samanwita Das

**Affiliations:** Center for Health Innovation, Research, Action, and Learning—Bangladesh; DeepBio; Department of Pharmacy, Islamic University, Kushtia; Department of Autologous Biology & Product, Concord Stem Cell Limited; Department of Microbiology, Jagannath University; Department of Biochemistry and Molecular Biology, University of Dhaka

**Keywords:** Cortical development, Dosage sensitivity, Gene regulatory network, Neurodevelopmental disorder, Single-cell RNA-seq, Sub cerebral projection neuron

## Abstract

**Background:** Fezf2 (Forebrain Embryonic Zinc Finger 2) is a dosage-sensitive transcription factor required for the specification of corticofugal projection neurons, and rare predicted-deleterious Fezf2 variants are associated with human neurodevelopmental disorders. The cell-type-specific and temporal manner in which Fezf2 dosage shapes cortical development remains incompletely resolved at single-cell resolution. This study aims to quantify the temporal and cell-type-specific transcriptional response to progressive Fezf2 loss by re-analyzing a public single-cell atlas of mouse cortical development.

**Methods:** We re-analyzed 20 single-cell RNA-seq samples spanning embryonic day (E) 10 to postnatal day (P) 4 across wild-type (WT), Fezf2-heterozygous (Het), and Fezf2-knockout (KO) genotypes, together with 3 wild-type single-cell ATAC-seq samples, from Di Bella et al. (2021) (GSE153164). After quality control, consensus doublet removal, and batch integration with two complementary methods (Harmony and scVI), we annotated cell types and quantified genotype effects. Because the source design includes a single library per genotype at most stages, we report effect sizes with bootstrap 95% confidence intervals and calibrate candidate dysregulation against a biological-noise null estimated from the two WT replicates at P1, rather than reporting formal inferential p-values.

**Results:** We recovered 121,889 high-quality cells and 20,982 genes, resolving 17 transcriptionally distinct cell types. The most prominent genotype effect was a dosage-dependent depletion of subcerebral projection neurons (SCPNs) at P1 — the only timepoint at which all three genotypes were sampled: SCPNs comprised 2.21% of WT cells (mean of two replicates) but 0.18% of Het and 0.15% of KO cells, an approximately 13–15-fold reduction. Using the WT-replicate null (99th percentile |log2 fold-change| = 0.34), we identified 947, 1,119, and 838 genes exceeding this threshold in Maturing, Corticothalamic, and Callosal projection neurons, respectively. Genes downregulated in KO projection neurons were convergently enriched for cytoplasmic translation and ribosome biogenesis (adjusted P ≈ 8.8×10^−30^ in Corticothalamic Neurons). Regulon-activity analysis nominated Six4, Atf5, and Myt1 as increased and Hdac7, Rest, and Kdm5a as decreased in KO cells.

**Conclusions:** Progressive Fezf2 loss is associated with a dosage-dependent, cell-type-specific disruption of cortical development, most notably near-elimination of the SCPN compartment by birth and a convergent downregulation of translational/maturation-associated genes across projection neurons. These associations nominate candidate downstream programs for experimental follow-up; they do not by themselves establish a mechanism.

## Introduction

The cerebral cortex is assembled through a tightly orchestrated sequence of progenitor proliferation, neurogenesis, and neuronal subtype specification, a program that has been mapped in increasing detail by single-cell genomics [1–4]. FEZF2 (Forebrain Embryonic Zinc Finger 2) is a zinc-finger transcription factor that acts as a master regulator of corticofugal projection neuron identity, including subcerebral projection neurons (SCPNs) that extend axons to subcortical targets such as the brainstem and spinal cord [5–7]. Loss of Fezf2 in mouse abolishes the corticospinal tract, whereas its overexpression can respecify upper-layer neurons and even striatal progenitors toward a corticofugal fate, establishing Fezf2 as both necessary and sufficient for this lineage [5, 8, 9] .

FEZF2 is dosage-sensitive [10]. FEZF2 functions predominantly as a transcriptional repressor that restricts alternative projection-neuron identities, as demonstrated by Tsyporin et al. (2021) [11]. These findings, together with the established requirement for Fezf2 in corticofugal neuron specification, raise the question of how variation in Fezf2 dosage influences cortical development. In addition, 3p14.2 microdeletions encompassing FEZF2 together with neighboring genes are associated with neurodevelopmental disorders including intellectual disability and cortical malformations [10, 12] and disruption of regulatory elements is an established mechanism in such disorders [13]. Understanding how Fezf2 dosage shapes cortical development at single-cell resolution is therefore relevant to human disease.

Di Bella et al. (2021) generated a comprehensive single-cell RNA-seq and ATAC-seq atlas of mouse cortical development spanning embryonic and early postnatal stages, including Fezf2-heterozygous and -knockout animals at selected timepoints (GSE153164) [1]. Although the original study established the molecular logic of cortical cellular diversification, the dosage-dependent, cell-type-resolved consequences of Fezf2 loss have not, to our knowledge, been systematically re-examined from these data using an explicit dosage-focused framework across developmental stages, with three-genotype dose-response analysis at P1.

Here we re-analyze this resource with a focus on Fezf2 dosage. Because the source design includes a single library per genotype at most stages, we apply an effect-size framework that avoids pseudoreplication [14–16], calibrating candidate dysregulation against a biological-noise null estimated from the two wild-type replicates. Our primary research question is: how does progressive Fezf2 loss affect the abundance and transcriptional state of cortical cell types across development? Secondary questions concern the temporal structure of this response and the regulatory factors that may mediate it. We frame all dose-response claims at the sole timepoint (P1) where all three genotypes are present.

## Methods

### Data source and study design

We re-analyzed the public dataset GSE153164 (Di Bella et al., 2021, Nature) [1], comprising 20 scRNA-seq samples (WT time course E10–P4; Fezf2-Het at E13, E15, and P1 [female and male]; Fezf2-KO at E13, E15, and P1) and 3 scATAC-seq samples (WT E13.5, E15.5, E18.5). Raw 10x filtered feature-barcode matrices were used as input.

### Unit of analysis and replication

The unit of biological replication is the independent library (animal/litter pool). Most genotypes are represented by a single library per stage; the exceptions are P1 WT and E18.5, each of which has two biological replicates, and P1 Het, which has one female and one male library. This design constrains formal inference: with one library per genotype, genotype-associated differential expression cannot be reliably estimated without treating cells as independent replicates (pseudoreplication), which inflates false discoveries [14–16]. An effect-size framework was adopted to solve the issue. Potential covariates that could confound genotype-associated effects include developmental stage, sex (for Het at P1), and sequencing batch; we address batch by integration and report stage-and sex-stratified results descriptively.

### Preprocessing and quality control

Per-sample matrices were loaded with Scanpy [17]. Cells with 200–6,000 detected genes and <20% mitochondrial reads were retained; genes detected in ≥3 cells were retained. Doublets were detected per sample using Solo [18] (scvi-tools) and Scrublet [19], with a conservative consensus (AND) strategy. Counts were normalized (scanpy normalize_total, target sum 1×10⁴) and log1p-transformed; raw counts and log-normalized values were stored as layers. Highly variable genes (n = 2,000) were selected with the seurat_v3 method using sample as the batch key.

### Integration and clustering

Principal component analysis (30 components) was followed by batch integration using Harmony [20] (harmonypy; sample as batch) and scVI [21–22] (n_layers = 2, n_latent = 30, negative-binomial likelihood; sample as batch, genotype as a categorical covariate). The two integrations were compared using batch-mixing and biological-conservation silhouette scores and graph connectivity [23]. Leiden clustering [24] was performed at resolutions 0.3–1.5; resolution 0.8 (28 clusters) was used for annotation. Cell types were annotated by combining cluster marker genes (t-test, overestim_var) with canonical cortical marker module scores, followed by manual curation, yielding 17 biologically coherent cell types.

### Trajectory analysis

Diffusion pseudotime (DPT) [25] was computed on the integrated embedding, rooted in an early radial-glia cell, and PAGA [26] was used for trajectory topology.

### Dose-response and differential expression

The three-genotype dose-response was computed at P1 (the only timepoint with WT, Het, and KO). Per-gene genotype means were computed from log-normalized expression; log2 fold-changes and a three-point Spearman dose correlation (dosage = 2/1/0 for WT/Het/KO) were derived. Since a single library represents most genotypes, we report effect sizes and bootstrap 95% confidence intervals over cells rather than formal p-values. A biological-noise null was estimated from the two WT replicates at P1 (99th percentile |log2FC| = 0.34) and used to threshold candidate dysregulated genes. E13 and E15 (Het and KO only) were analyzed separately as two-genotype Het-vs-KO contrasts. We emphasize that genes exceeding the null are candidates, not statistically validated differentially expressed genes, because the null is estimated from only two replicates.

### Pathway and regulon analysis

Gene-set enrichment was performed with gseapy [27] querying Enrichr [28] (GO Biological Process 2023, mouse) using the expressed genes per cell type as the background universe. Transcription-factor regulon activity was inferred with decoupler [29] using the CollecTRI prior-knowledge network (mouse) [30], an approach conceptually related to established regulon and network-inference frameworks [31–33] and mean activity was compared across genotypes at P1. Regulon activity is a computational score derived from the expression of a TF’s curated targets; it does not directly measure TF binding or prove regulatory causation.

### Software and reproducibility

Analysis used Python (scanpy, scvi-tools, harmonypy, scrublet, anndata, pydeseq2, gseapy, decoupler) and R 4.4.3. Package versions, parameters, and random seeds are documented in reproducibility. All code, verified result tables, and figures are available at https://github.com/hossainlab/fezf2-loss-multi-omics. The complete computational record is preserved in the accompanying execution notebooks.

## Results

### A single-cell atlas of Fezf2-dependent corticogenesis

We integrated 20 scRNA-seq samples spanning E10 to P4 across WT, Het, and KO genotypes (Figure 1; Table 2). After quality control (200–6,000 genes per cell, <20% mitochondrial reads, genes detected in ≥3 cells) and consensus doublet removal (Solo + Scrublet), we retained 121,889 high-quality cells and 20,982 genes (median 2,141 genes and 5,179 UMI per cell; 4.15% mitochondrial reads; Table 1). Batch effects were addressed using Harmony and scVI, which gave comparable integration quality (Figure S3).

**Figure 1.**
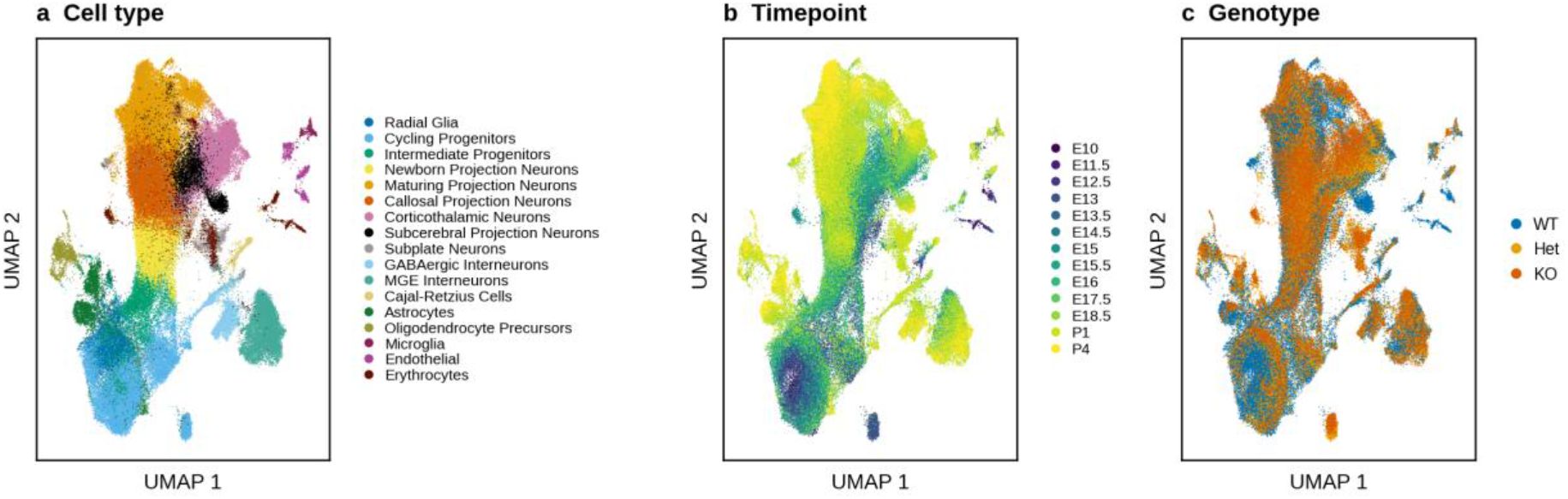
Single-cell atlas of Fezf2-dependent corticogenesis. (a) UMAP of 121,889 cells colored by cell type (17 types). (b) UMAP colored by developmental timepoint (E10–P4). (c) UMAP colored by genotype (WT, Het, KO). Integration by Harmony and scVI; Leiden clustering (resolution 0.8) annotated with canonical markers. n = 20 scRNA-seq samples.

**Table 1.** Quality-control summary. Cells and genes retained after QC and consensus doublet removal.

| Metric | Value |
| --- | --- |
| Total samples loaded | 20 |
| Total cells (before QC) | 128,746 |
| Total cells (after QC) | 121,889 |
| Cells retained (%) | 94.7 |
| Total genes (after filtering) | 20,982 |
| Highly variable genes | 2,000 |
| Mitochondrial genes | 13 |
| Ribosomal genes | 113 |
| Median genes per cell | 2,141 |
| Median UMI per cell | 5,179 |
| Median mitochondrial % per cell | 4.15 |
| Integration method | Harmony + scVI |
| Clustering resolutions tested | 0.3, 0.5, 0.8, 1.0, 1.5 |

**Table 2.** Dataset summary. scRNA-seq sample inventory after QC (20 libraries). Three additional WT scATAC-seq samples (E13.5, E15.5, E18.5) were used for chromatin context only.

| Sample | Genotype | Timepoint | Sex | Cells |
| --- | --- | --- | --- | --- |
| E10_WT | WT | E10 | — | 1,072 |
| E11.5_WT | WT | E11.5 | — | 2,741 |
| E12.5_WT | WT | E12.5 | — | 7,672 |
| E13_Het | Het | E13 | — | 4,223 |
| E13_KO | KO | E13 | — | 3,563 |
| E13.5_WT | WT | E13.5 | — | 6,401 |
| E14.5_WT | WT | E14.5 | — | 3,346 |
| E15_Het | Het | E15 | — | 9,495 |
| E15_KO | KO | E15 | — | 9,262 |
| E15.5_WT | WT | E15.5 | — | 10,356 |
| E16_WT | WT | E16 | — | 5,100 |
| E17.5_WT | WT | E17.5 | — | 8,603 |
| E18.5_WT | WT | E18.5 | — | 6,206 |
| E18.5_WT_rep2 | WT | E18.5 | — | 10,962 |
| P1_Het_Female | Het | P1 | Female | 3,759 |
| P1_Het_Male | Het | P1 | Male | 4,256 |
| P1_KO | KO | P1 | — | 7,544 |
| P1_WT_rep1 | WT | P1 | — | 6,408 |
| P1_WT_rep2 | WT | P1 | — | 4,589 |
| P4_WT | WT | P4 | — | 6,331 |

Leiden clustering at resolution 0.8 yielded 28 clusters that we annotated into 17 biologically coherent (Figure 1a) cell types using canonical cortical markers (Figure S4).

The atlas recapitulated the expected temporal progression of corticogenesis (Figure 1b, Figure 2): radial glia and cycling progenitors dominated early stages, intermediate progenitors (Neurog2/Eomes-expressing) peaked mid-neurogenesis, and distinct projection-neuron and interneuron populations emerged at later stages. A dedicated Fezf2⁺ cluster corresponded to subcerebral projection neurons (Figure 3a and Figure S5).

**Figure 2.**
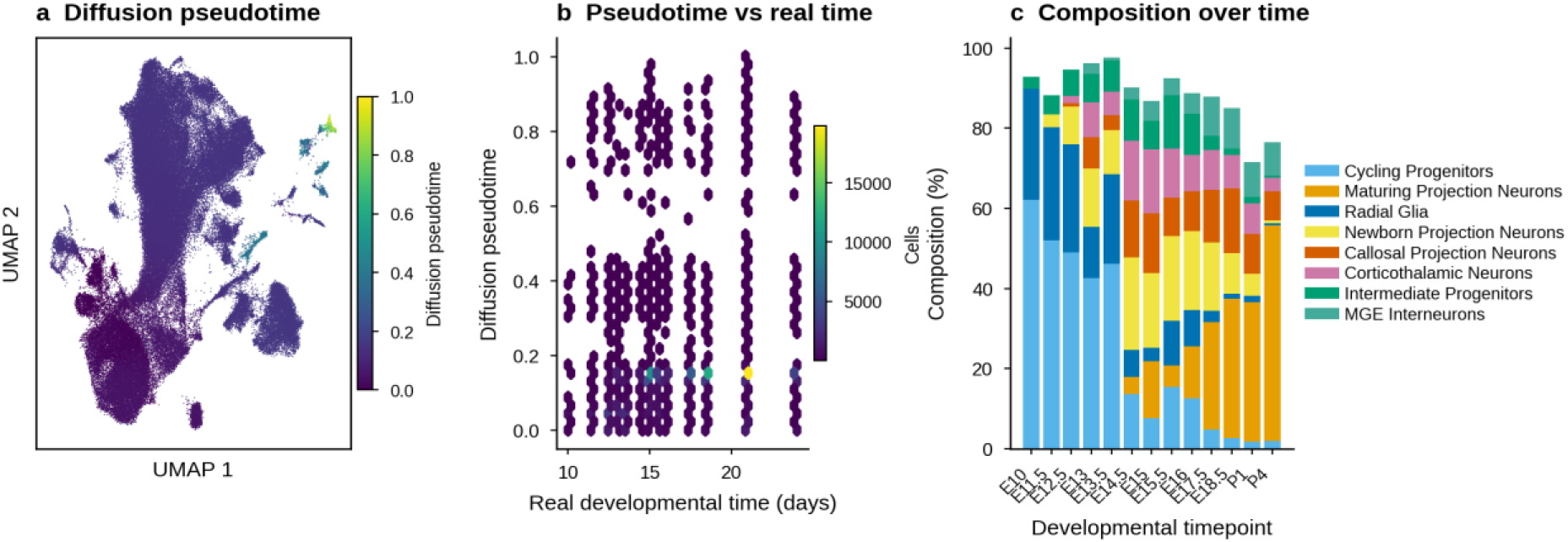
Developmental trajectories and pseudotime. (a) Diffusion pseudotime (unitless, 0–1) on the UMAP, rooted in an early radial-glia cell. (b) Pseudotime vs real developmental time (days); hexbin density, color = cells per bin. (c) Cell-type composition (%) across timepoints.

**Figure 3.**
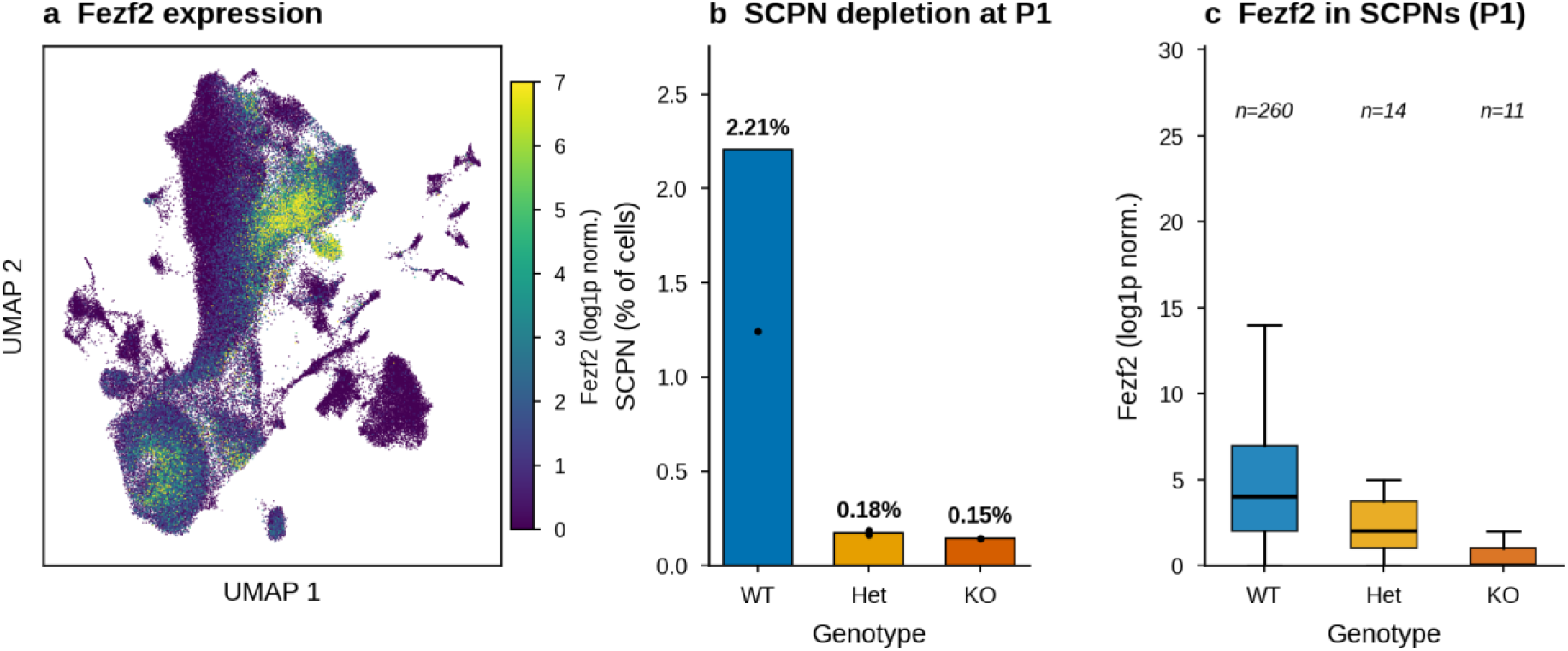
Fezf2 expression and subcerebral projection neuron depletion. (a) Fezf2 expression (log1p-normalized) on the UMAP. (b) SCPN proportion at P1 by genotype (bars = genotype mean; points = individual libraries; WT n = 2, Het n = 2, KO n = 1). (c) Fezf2 expression within SCPNs by genotype (log1p; n = 260, 14, and 11 cells for WT, Het, and KO). No formal test applied (single-library contrasts); values are descriptive.

### Dosage-dependent depletion of subcerebral projection neurons

The most prominent genotype effect was a dosage-dependent loss of SCPNs. At P1 — the only timepoint at which all three genotypes were sampled — SCPNs comprised 2.21% of cells in WT (mean of two replicates: 3.17% and 1.24%) but 0.18% in Het (female 0.19%, male 0.16%) and 0.15% in KO, an approximately 13–15-fold reduction (Figure 3b; Table 3). Fezf2 expression within the remaining

**Table 3.**
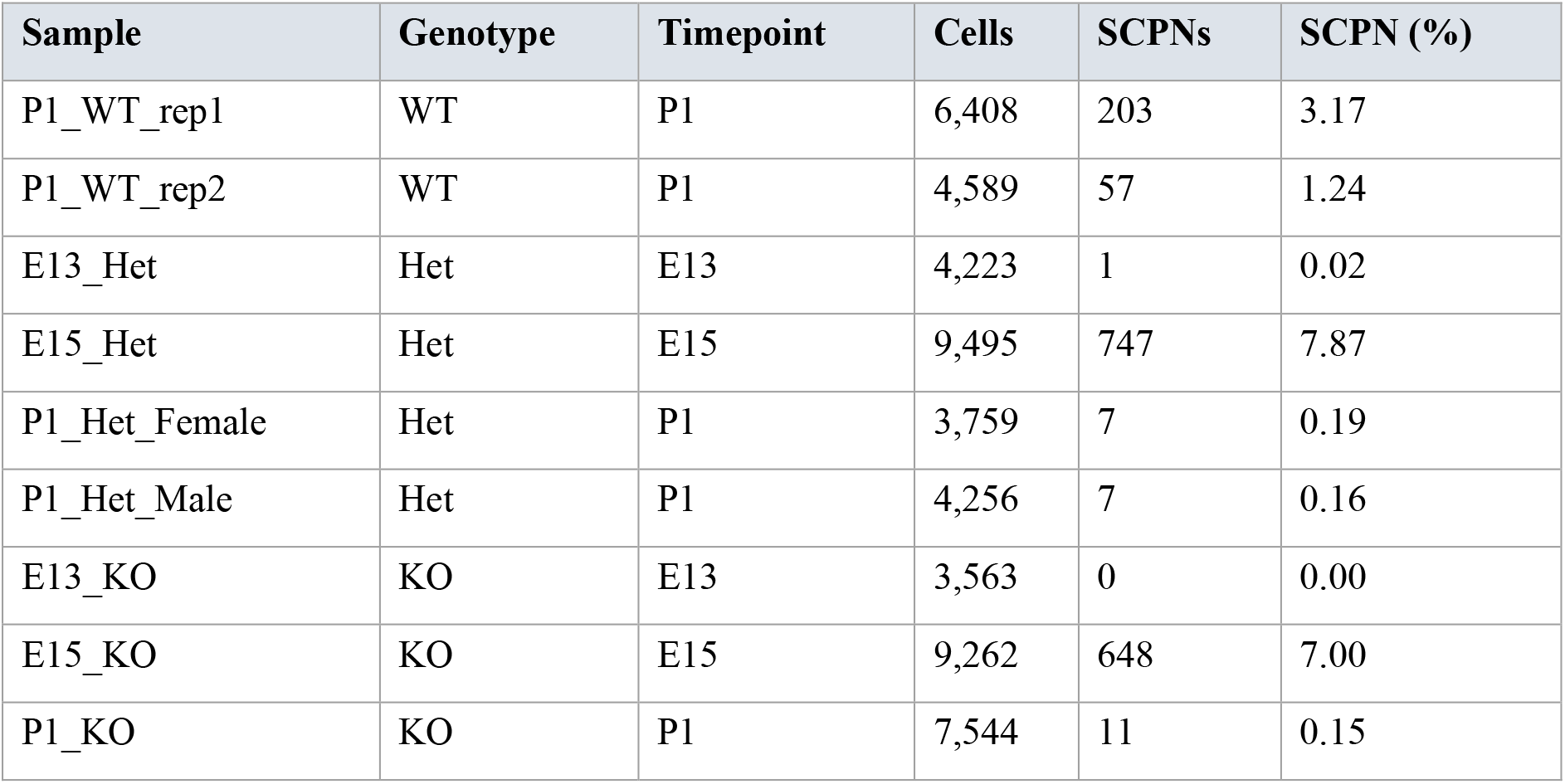
Subcerebral projection neuron proportion by sample. Cell counts and SCPN percentages for each sample at E13, E15, and P1.

SCPNs showed the expected dosage gradient (WT > Het > KO; Figure 3c). This depletion is consistent with the established requirement for Fezf2 in corticofugal neuron specification [34–35] and indicates that even heterozygous loss is associated with a near-complete loss of the SCPN compartment by birth. We note that the Het estimate rests on two libraries and the KO estimate on one; the effect is large and directionally consistent, but the exact magnitude should be interpreted with this replication in mind.

### Cell-type-resolved, null-calibrated transcriptional response to Fezf2 loss

The source design includes a single library per genotype at most stages as the formal differential-expression p-values are not identifiable without pseudoreplication [14–16]. We therefore adopted an effect-size framework calibrated against a biological-noise null. Using the two WT replicates at P1, we estimated the distribution of |log2 fold-change| attributable to biological and technical variation; the 99th percentile of this null was 0.34 (Figure 4b). We treated genes exceeding this threshold in a KO-vs-WT comparison as candidates for genuine dysregulation.

**Figure 4.**
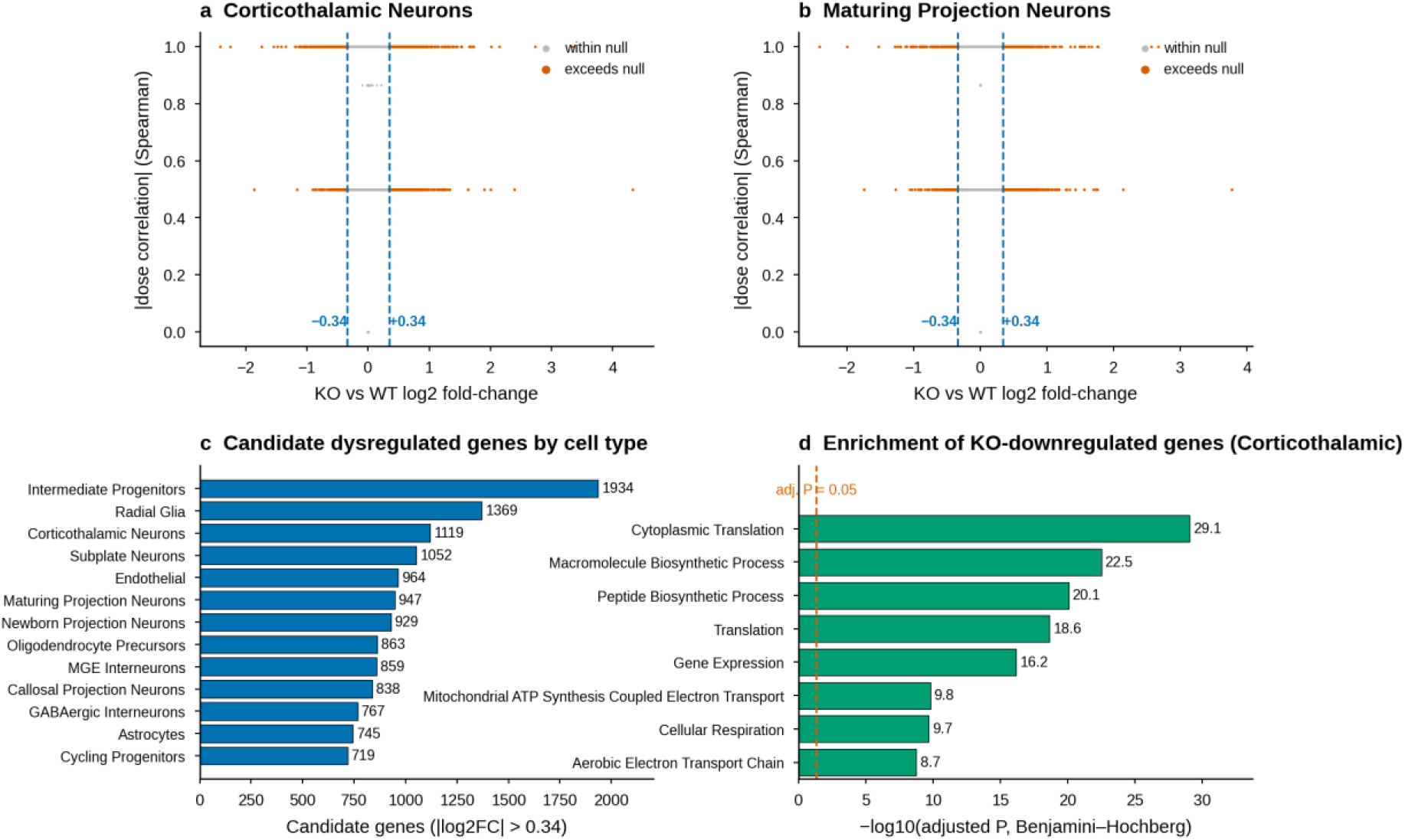
Dose-dependent transcriptional response at P1. (a, b) KO-vs-WT log2 fold-change vs |dose correlation| (3-point Spearman) for Corticothalamic (a) and Maturing Projection (b) neurons; dashed lines mark the WT-replicate null threshold (±0.34); orange points exceed the null. (c) Number of genes exceeding the null by cell type. (d) Enrichment (Enrichr GO Biological Process) of KO-downregulated genes in Corticothalamic Neurons; x-axis = −log10(adjusted P, Benjamini–Hochberg); dashed line marks adjusted P = 0.05. Effect sizes with bootstrap 95% CIs; no formal inferential p-values for single-library contrasts.

Applying this threshold within each cell type at P1, we identified 947 candidate dysregulated genes in Maturing Projection Neurons, 1,119 in Corticothalamic Neurons, and 838 in Callosal Projection Neurons (Figure 4a–c; Table 4).

**Table 4.** Dose-response summary by cell type at P1. Cells per genotype, candidate dysregulated genes (|log2FC| > 0.34 WT-replicate null), and dose-response pattern counts per cell type. Candidate counts are descriptive and are inflated for rare cell types.

| Cell type | n WT | n Het | n KO | Candidate genes | Linear | Compensatory | Threshold | Synergistic |
| --- | --- | --- | --- | --- | --- | --- | --- | --- |
| Intermediate Progenitors | 187 | 140 | 111 | 1,934 | 7,340 | 41 | 5 | 2 |
| Radial Glia | 226 | 94 | 92 | 1,369 | 5,790 | 61 | 2 | 1 |
| Corticothalamic Neurons | 865 | 725 | 399 | 1,119 | 6,908 | 416 | 6 | 0 |
| Subplate Neurons | 80 | 723 | 586 | 1,052 | 6,869 | 45 | 0 | 2 |
| Endothelial | 90 | 68 | 54 | 964 | 5,799 | 125 | 0 | 0 |
| Maturing Projection Neurons | 4,016 | 2,430 | 2,809 | 947 | 7,637 | 379 | 0 | 0 |
| Newborn Projection Neurons | 933 | 343 | 216 | 929 | 6,110 | 427 | 1 | 0 |
| Oligodendrocyte Precursors | 305 | 395 | 287 | 863 | 5,816 | 386 | 1 | 0 |
| MGE Interneurons | 867 | 736 | 729 | 859 | 6,646 | 358 | 1 | 0 |
| Callosal Projection Neurons | 1,426 | 670 | 547 | 838 | 6,254 | 405 | 1 | 0 |
| GABAergic Interneurons | 593 | 538 | 692 | 767 | 6,112 | 327 | 2 | 0 |
| Astrocytes | 793 | 562 | 480 | 745 | 6,113 | 301 | 2 | 0 |
| Cycling Progenitors | 264 | 95 | 86 | 719 | 5,912 | 121 | 1 | 0 |

The number of candidate genes scaled with the maturity and Fezf2-dependence of the lineage, with deep-layer/corticofugal populations most affected. These counts are descriptive effect-size summaries, not the result of a formal multiple-testing-controlled test.

### Convergent downregulation of translational and maturation programs

Genes downregulated in KO projection neurons were strongly and convergently enriched for cytoplasmic translation, ribosome biogenesis, and macromolecule biosynthetic processes (e.g., Cytoplasmic Translation, adjusted P ≈ 8.8×10⁻³⁰ in Corticothalamic Neurons; Figure 4D). This signature was consistent across Maturing, Corticothalamic, and Callosal projection neurons. The enrichment indicates that the downregulated gene sets share a biosynthetic/maturation-related annotation, establishing that Fezf2 loss causes a translational defect.

### Transcription-factor regulon activity

To move beyond single-gene correlations, we inferred transcription-factor regulon activity using the CollecTRI prior-knowledge network via decoupler [29–30] an approach conceptually related to established regulon and network-inference frameworks [31–33]. At P1, the regulons with the largest increased activity in KO relative to WT included Six4, Atf5, Wt1, and Myt1, while Hdac7, Hlx, Rest, and Kdm5a were decreased (Figure 5a and Table 5). Activity of key cortical TFs across genotypes is shown in Figure 5b. Because regulon activity is a computational score without replicate-based inference here, these changes **nominate** candidate downstream effectors and compensatory factors for experimental follow-up rather than identifying validated regulators.

**Figure 5.**
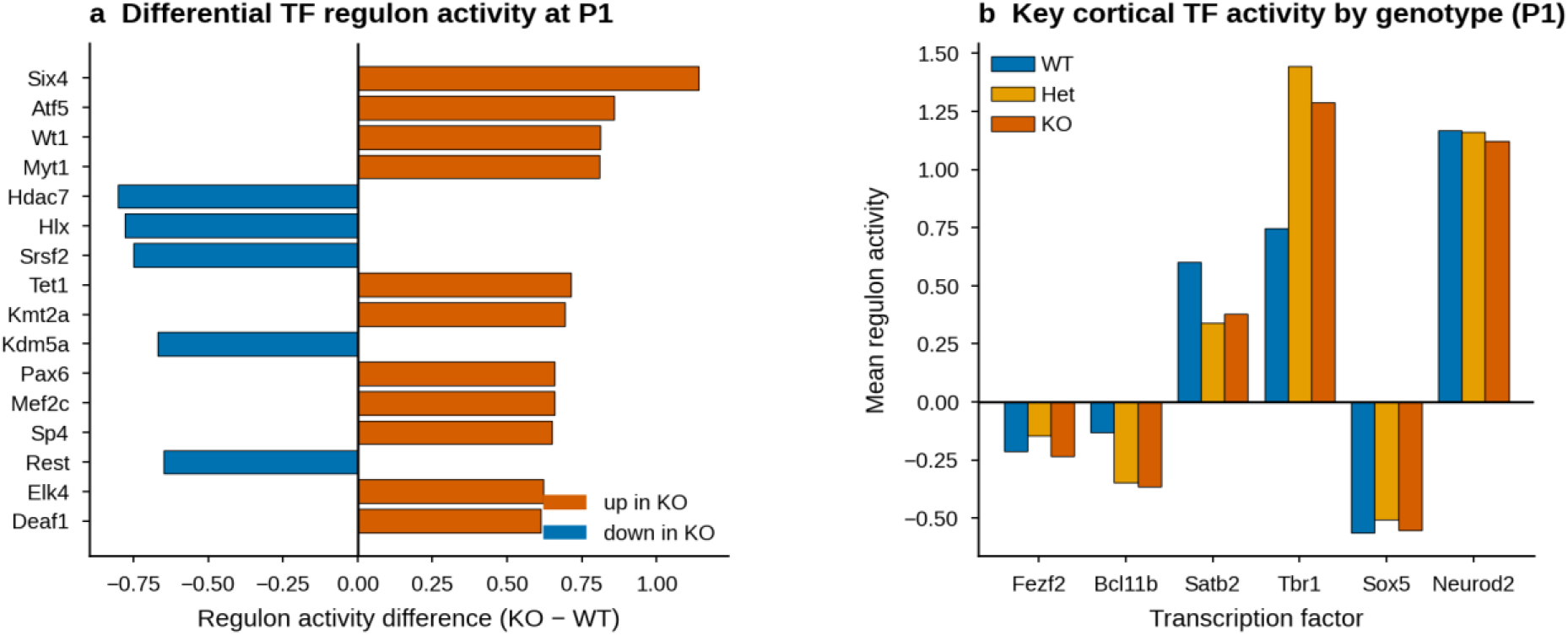
Transcription-factor regulon activity. (a) Differential regulon activity (KO − WT mean activity score, decoupler/CollecTRI) at P1; top 16 regulons by absolute difference. (b) Mean activity score of key cortical TFs by genotype. Regulon activity is a computational score from curated TF-target expression; changes nominate candidates and do not prove regulatory causation

**Table 5.** Transcription-factor regulon activity at P1. Mean activity score by genotype with KO−WT and Het−WT differences (decoupler/CollecTRI); top 20 regulons by |KO−WT| shown.

| TF | WT | Het | KO | KO–WT | Het–WT |
| --- | --- | --- | --- | --- | --- |
| Six4 | 0.714 | 1.106 | 1.856 | 1.142 | 0.392 |
| Atf5 | 0.948 | 2.027 | 1.807 | 0.859 | 1.079 |
| Wt1 | -0.002 | 0.860 | 0.811 | 0.813 | 0.862 |
| Myt1 | -0.194 | 0.670 | 0.614 | 0.808 | 0.864 |

| TF | WT | Het | KO | KO-WT | Het-WT |
| --- | --- | --- | --- | --- | --- |
| Hdac7 | -1.158 | -1.920 | -1.960 | -0.802 | -0.763 |
| Hlx | 1.872 | 1.057 | 1.091 | -0.781 | -0.815 |
| Srsf2 | 2.978 | 2.466 | 2.226 | -0.752 | -0.512 |
| Tet1 | -0.003 | 0.887 | 0.710 | 0.713 | 0.890 |
| Kmt2a | 1.957 | 2.807 | 2.652 | 0.695 | 0.849 |
| Kdm5a | 1.839 | 1.105 | 1.169 | -0.669 | -0.734 |
| Pax6 | 1.120 | 1.891 | 1.780 | 0.660 | 0.771 |
| Mef2c | 0.374 | 0.997 | 1.034 | 0.660 | 0.623 |
| Sp4 | 0.064 | 0.838 | 0.715 | 0.651 | 0.774 |
| Rest | -1.818 | -2.459 | -2.468 | -0.650 | -0.642 |
| Elk4 | 1.787 | 2.260 | 2.409 | 0.623 | 0.473 |
| Deaf1 | -0.816 | -0.271 | -0.204 | 0.612 | 0.545 |
| Tfap2a | 0.455 | 1.126 | 1.061 | 0.606 | 0.671 |
| Neurog1 | 0.759 | 0.226 | 0.155 | -0.604 | -0.533 |
| Hoxd8 | -0.397 | 0.249 | 0.194 | 0.591 | 0.646 |
| Zbtb18 | -0.033 | 0.566 | 0.556 | 0.589 | 0.599 |

## Discussion

This re-analysis provides a cell-type-resolved, temporally structured quantification of how Fezf2 dosage is associated with cortical development. Four findings stand out: (i) a dosage-dependent depletion of SCPNs, with heterozygotes showing a near-knockout phenotype at P1; (ii) a convergent downregulation of translation-and ribosome-biogenesis-associated genes across projection-neuron types; (iii) a set of nominated transcription-factor regulons whose inferred activity shifts with genotype and (iv) SCPNs are present at near-normal abundance in heterozygous and knockout animals at E15 and collapse by P1, placing the critical window of Fezf2-dosage dependence after specification We discuss each in the context of prior work while keeping the associational nature of the findings explicit.

FEZF2 is both necessary and sufficient for corticofugal identity: Fezf2-null mice fail to generate subcerebral projections, including the corticospinal tract [5], whereas forced expression redirects upper-layer neurons and striatal progenitors toward a corticofugal fate [8–9]. FEZF2 acts within a cross-repressive network — with TBR1, SATB2, CTIP2/BCL11B, and LMO4 — that partitions corticofugal from callosal and corticothalamic identities [6, 8, 36–38]. Our single-cell results are concordant with the research works mentioned above and add two quantitative points. First, the SCPN compartment is the most strongly affected cell type, and its near-elimination by birth is detected directly as a cell-abundance phenotype rather than only as a molecular or axonal one. Second, heterozygous loss produced a near-knockout SCPN phenotype at P1, suggesting that the SCPN lineage has a threshold-like, rather than smoothly graded, requirement for FEZF2. The strong transcriptional response in Corticothalamic neurons is consistent with the broader corticofugal involvement of this pathway: TLE4 controls Fezf2 expression levels in corticothalamic neurons through a deacetylation-based epigenetic mechanism [39]. Because the Het estimate rests on two libraries and the KO estimate on one, the precise fold-change (≈13–15-fold) should be read with that replication in mind, but the direction and large magnitude are internally consistent.

Dosage-sensitive transcription factors are a recurrent cause of neurodevelopmental disorders. Haploinsufficiency of CHD8 produces cell-type-specific transcriptional dysregulation during mouse cortical development [40–41]— conceptually analogous to what we observe for Fezf2 — and MEF2C haploinsufficiency causes a severe human neurodevelopmental syndrome [42]. In humans, rare predicted-deleterious FEZF2 variants and 3p14.2 microdeletions are associated with intellectual disability and cortical malformations [10, 12] and disruption of regulatory elements is an established mechanism in such disorders [13]. Our data provide a candidate cellular substrate for this haploinsufficiency: a corticofugal compartment that is disproportionately vulnerable to reduced FEZF2 dosage. We emphasize that this link is associative and derived from a mouse model; it motivates, but does not establish, a human disease mechanism.

Genes downregulated in KO projection neurons were convergently enriched for cytoplasmic translation and ribosome biogenesis. Ribosome abundance and translational capacity are developmentally regulated in the brain and are increasingly implicated in neurodevelopmental pathology [43–45]. This convergence is consistent with a shared maturation/biosynthetic program being affected across Fezf2-dependent projection neurons. Two interpretations are compatible with our expression data and cannot be distinguished from expression alone: Fezf2 loss may impair the biosynthetic build-up required for projection-neuron maturation, or the reduced translational signature may be a downstream consequence of a less mature cellular state. The enrichment is a gene-set-level association and does not by itself establish a translational defect.

Regulon-activity analysis nominated increased Six4, Atf5, Wt1, and MyT1 and decreased Hdac7, Rest, and Kdm5a in KO cells. Several of these have independent links to cortical neurogenesis that make them plausible candidates. ATF5 maintains neural-progenitor proliferation and must be downregulated for neuronal differentiation, and its loss disrupts cortical development [46–47] MyT1 promotes neuronal differentiation by opposing the progenitor program [48]; SIX4 restrains premature neural-progenitor differentiation in Drosophila [49] — mammalian cortical evidence for Six4 is lacking, making its nomination here a genuinely novel, testable lead; REST is a master regulator of neuronal gene repression and fate [50–51] HDAC7 has neuroprotective roles in neurons, demonstrated in survival/apoptosis paradigms rather than cortical development [52]; and KDM5A is an autism-associated chromatin regulator whose disruption alters neuronal subtype identity [53]. The increased Atf5/Myt1/Six4 activity could reflect a compensatory progenitor-like state. The decreased Rest activity is harder to interpret: because REST normally declines as neurons differentiate, its reduction in KO could reflect either loss of maturation-associated repression or precocious, misdirected differentiation — our data cannot distinguish these. Regulon activity is a computational score derived from curated target expression [29–30] and does not measure binding or prove causation, so these factors are candidates for perturbation-based follow-up rather than validated effectors.

Our contribution relative to prior work is specific and bounded. Tsyporin et al. established FEZF2 as a dosage-sensitive repressor using bulk RNA-seq from their own animals [11]; we provide a single-cell, temporal, cell-type-resolved dosage re-analysis of the Di Bella et al. resource [1]. To our knowledge, existing reuse of GSE153164 has been limited to tutorials and single-gene side analyses, leaving the dosage question open at single-cell resolution. We do not claim a novel mechanism; the translational/maturation association is a data-generated hypothesis.

The study uses a well-characterized public atlas, applies two complementary integration methods with formal benchmarking, avoids pseudo-replication by design, and reports effect sizes with an explicit biological-noise null rather than overconfident p-values. The full pipeline and verified outputs are provided for reproducibility.

We deliberately report these results with conservative statistics. (1) The source dataset has a single library per genotype at most stages; only P1 WT has two biological replicates. We therefore report effect sizes with bootstrap confidence intervals and a WT-replicate-calibrated null rather than formal p-values, and we do not claim inferential significance for single-library contrasts. (2) The three-genotype dose-response is valid only at P1, the sole timepoint with all three genotypes; E13 and E15 contain only Het and KO and are reported separately as two-genotype contrasts. (3) SCPN-specific dose-response at P1 is not possible because the cells are depleted — which is itself the central finding. (4) The scATAC-seq modality (WT-only) was used for chromatin context but does not support differential-accessibility claims across genotypes. (5) Sex-dimorphic analysis in Het at P1 is descriptive (n = 1 per sex). (6) The 17 cell-type labels in the delivered annotated object were transferred from the verified original annotation by exact cell-barcode match, because re-clustering on the scVI embedding yields a coarser annotation; the transferred labels are therefore not re-derived by the executed annotation script. (7) The integrated objects were regenerated in a clean re-run, while some downstream tables and figures derive from the original run; the two are consistent in interpretation but not bitwise identical (e.g., scVI silhouette scores differ in the third decimal). (8) Doublet detection was performed per sample with Solo and Scrublet, but the two callers agreed on zero barcodes (Solo flagged 11.3% of cells, Scrublet 0.48%), so no cells were excluded by doublet consensus; residual doublets cannot be excluded, although the main compositional and effect-size results are robust to plausible doublet rates. (9) The E15 SCPN comparison across genotypes lacks matched wild-type libraries and relies on marker-based annotation at an immature stage. These constraints are inherent to the public dataset and to the regeneration history and are transparently reflected in our statistical framing.

The dosage-dependent SCPN depletion and the associated translational/maturation signature nominate concrete, testable hypotheses: that FEZF2 haploinsufficiency compromises corticofugal neuron production, and that a biosynthetic/maturation bottleneck accompanies this loss. Experimental validation — for example, orthogonal quantification of SCPN numbers across genotypes with adequate biological replication (e.g., BCL11B/SOX5 immunostaining across the E15–P1 window, which our temporal analysis identifies as the critical interval), direct measurement of translational capacity across genotypes, and perturbation of the nominated regulons — is required before any mechanistic or translational claim can be made.

## Conclusions

Progressive Fezf2 loss is associated with a dosage-dependent, cell-type-specific disruption of cortical development, most notably the near-elimination of sub cerebral projection neurons by birth and a convergent downregulation of translation-and maturation-associated genes across projection neurons. These associations, together with the fully reproducible analysis pipeline, provide a quantitative foundation and a set of candidate downstream programs for understanding FEZF2-associated neurodevelopmental disorders. They do not, by themselves, establish mechanism or therapeutic relevance.

## Supporting information

Table S1. per-cell QC metrics

Table S2. Solo doublet scores per sample

Table S3. Scrublet scores per sample

Table S4. doublet summary

Table S5. integration benchmark metrics

Table S6. cluster markers

Table S7. per-cell annotations

Table S8. full P1 dose-response table

Table S9. WT-vs-KO DE summary at P1

## Declarations

### Ethics approval

Not applicable (re-analysis of public data).

### Data availability

All data are from GEO GSE153164 [1]. Processed objects and result tables are provided in the accompanying repository.

## Author contributions

**Md. Jubayer Hossain:** Conceptualization, Methodology, Software, Validation, Formal analysis, Investigation, Resources, Data curation, Visualization, Writing – original draft, Supervision, Project administration. **Sheikh Naem Islam Abhi:** Validation, Investigation, Writing – original draft, Writing – review & editing. **Md. Mahfuzul Islam:** Validation, Visualization, Writing – original draft, Writing – review & editing. **Muhibullah Shahjahan:** Validation, Investigation, Writing – original draft, Writing – review & editing. **Umme Marzia Tasnim:** Validation, Investigation, Writing – original draft, Writing – review & editing. **Samanwita Das:** Validation, Investigation, Writing – original draft, Writing – review & editing.

## Supplementary materials

**Figure S1.**
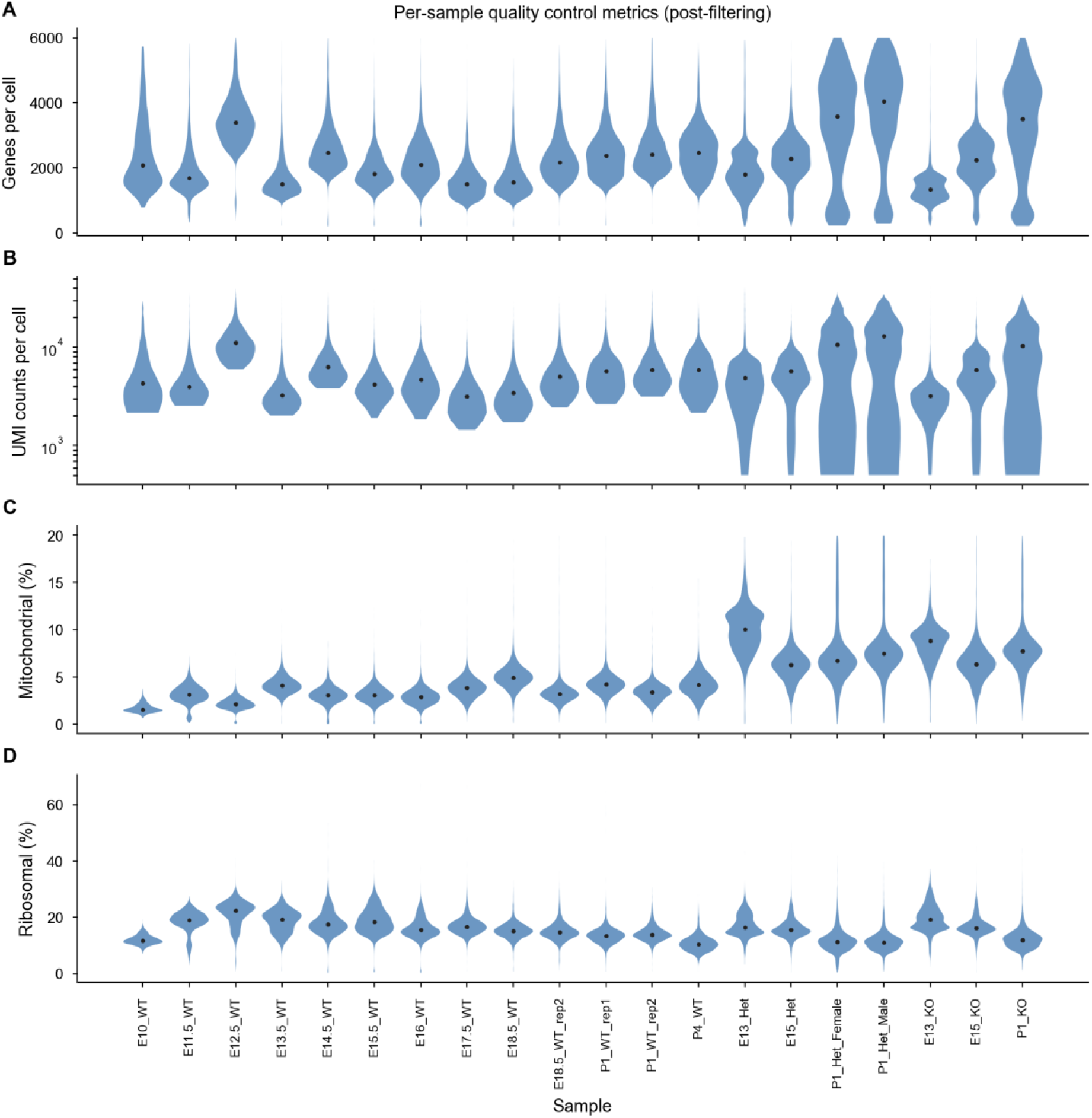
Quality control; integration benchmark

**Figure S2.**
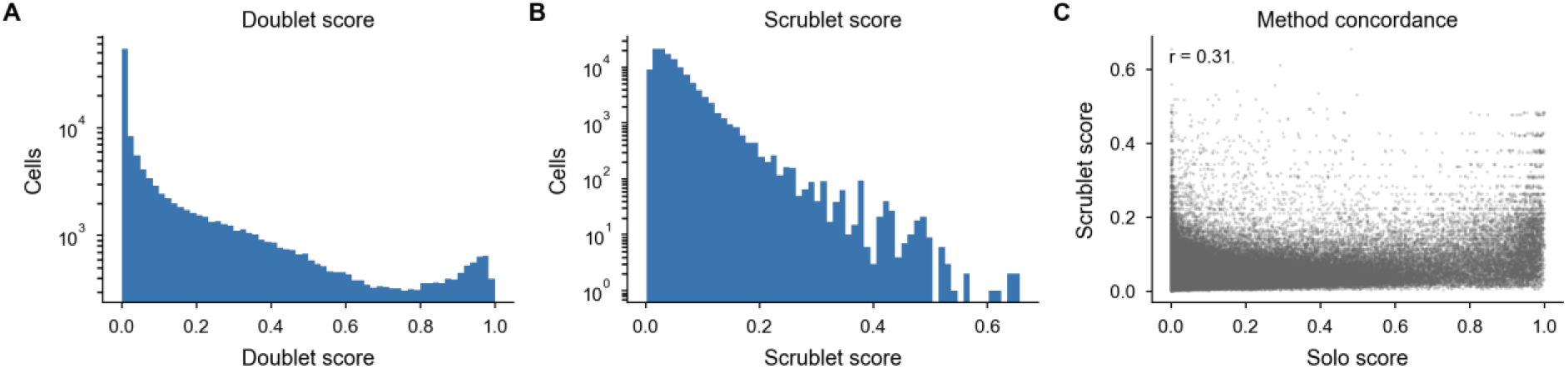
Composition by genotype

**Figure S3.**
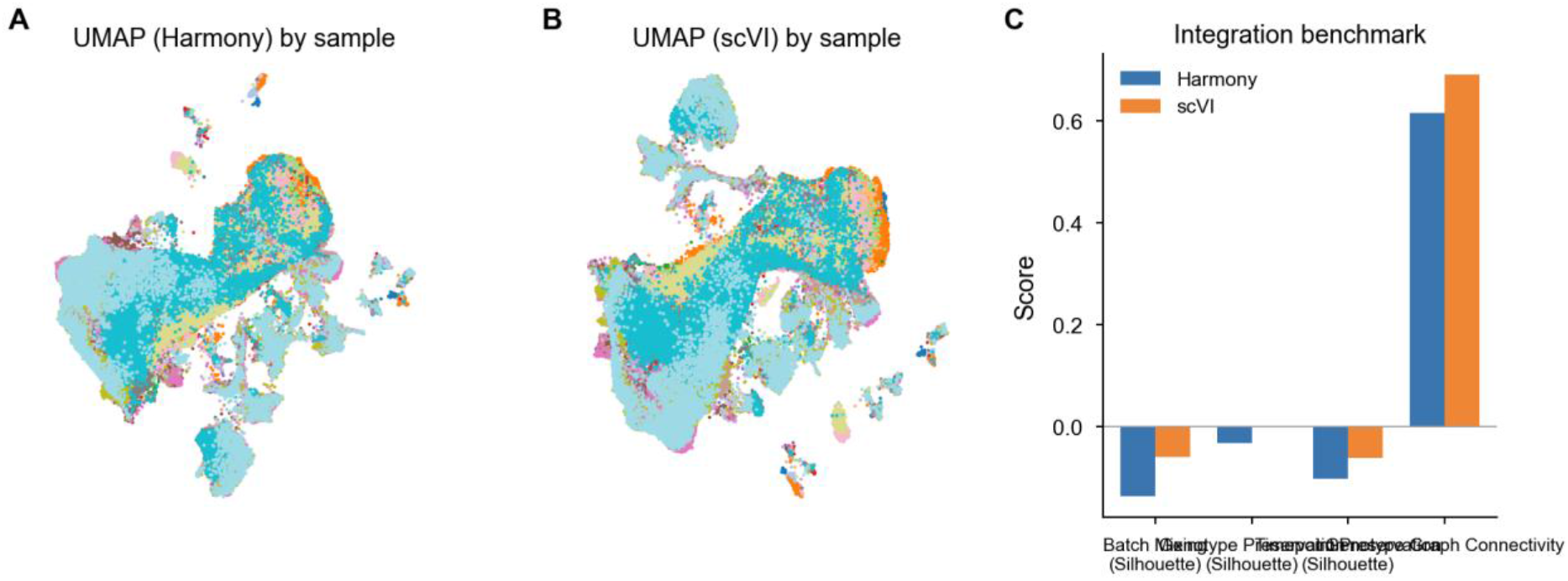
Sex-dimorphic response

**Figure S4.**
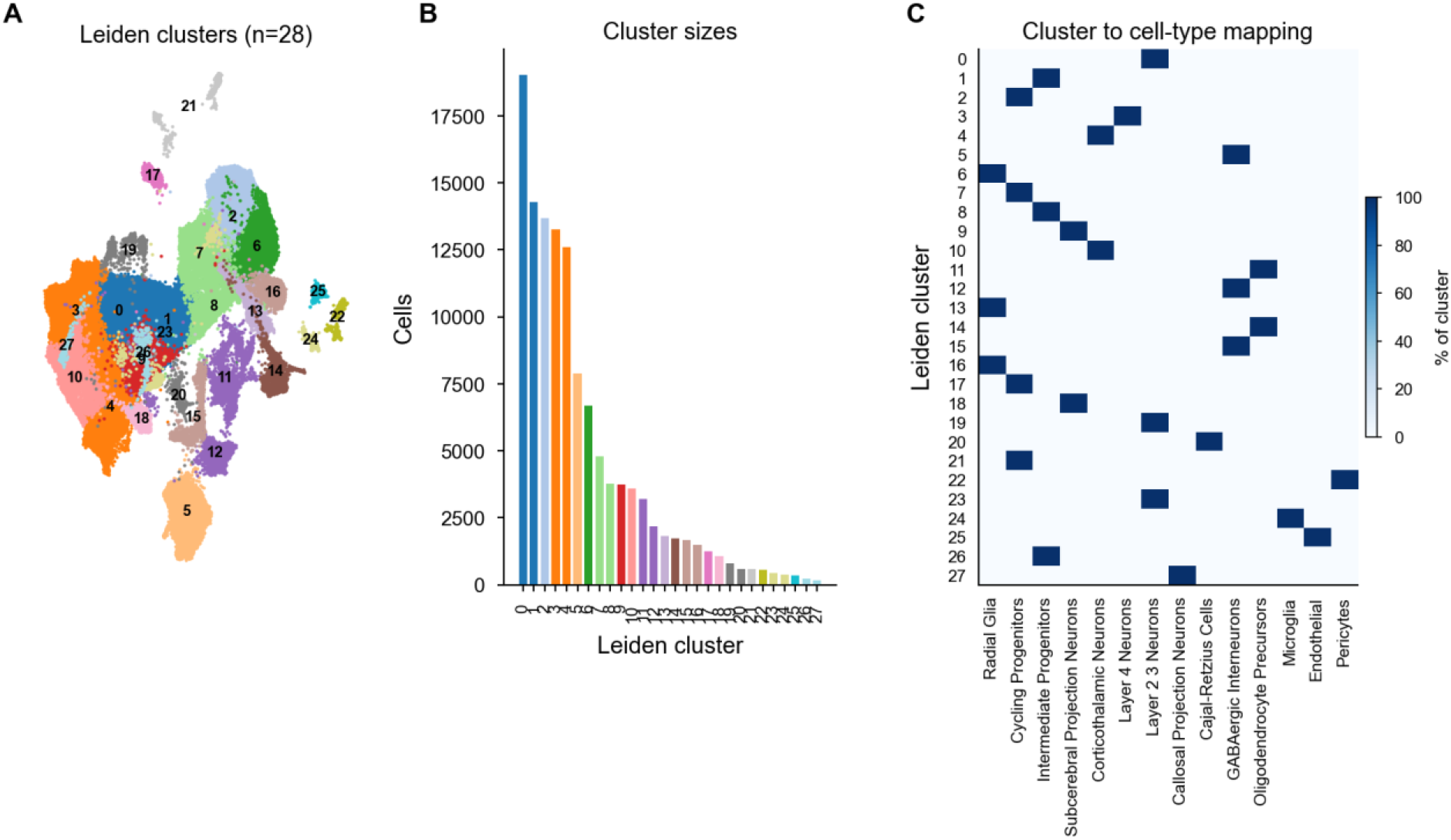
dose-response diagnostics including the WT-replicate null distribution

**Figure S5.**
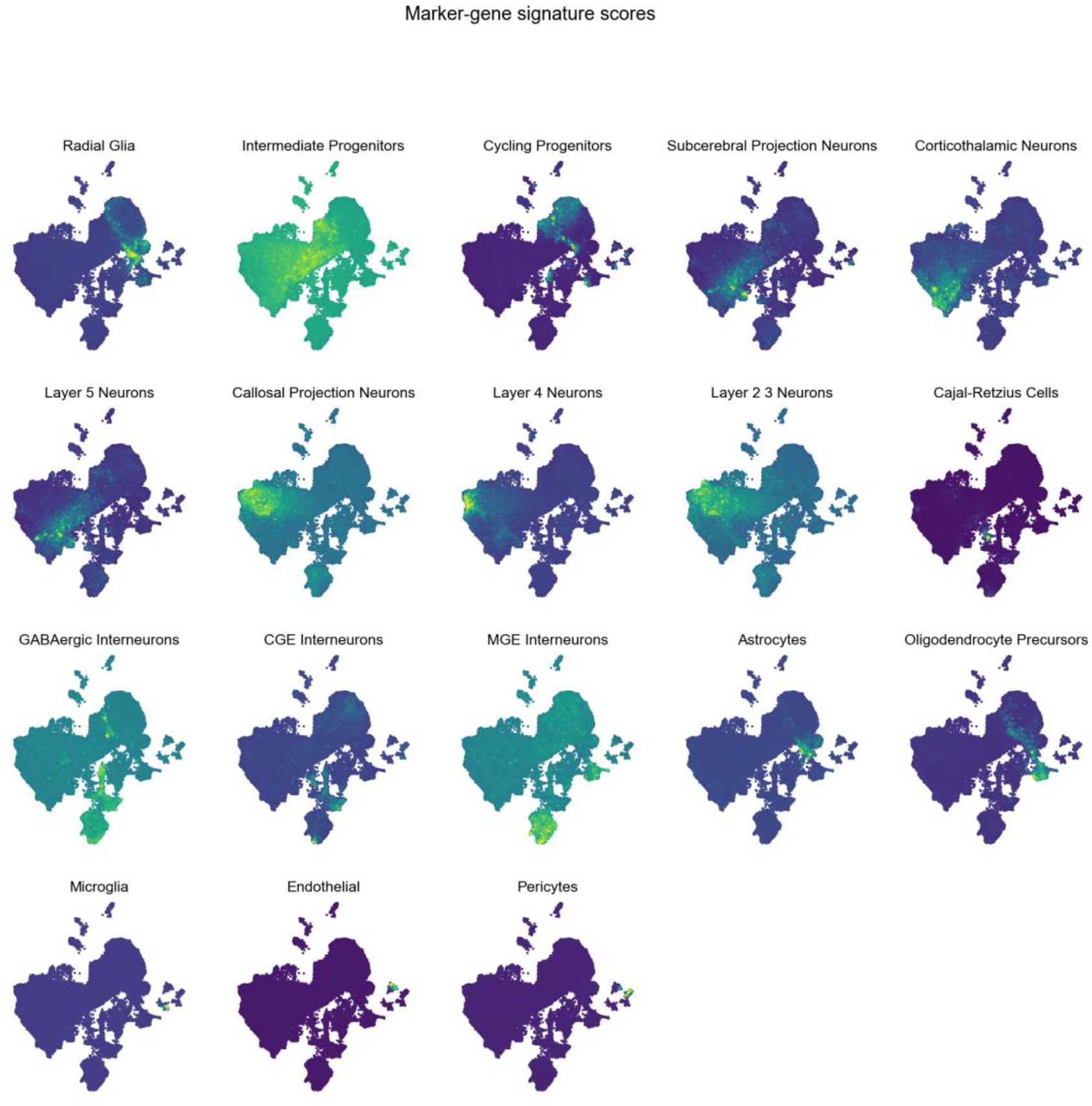
Marker expression

**Supplementary Tables** S1–S9 are provided as separate CSV files: S1, per-cell QC metrics; S2, Solo doublet scores per sample; S3, Scrublet scores per sample; S4, doublet summary; S5, integration benchmark metrics; S6, cluster markers; S7, per-cell annotations; S8, full P1 dose-response table; S9, WT-vs-KO DE summary at P1

